# DOMON domain-containing proteins control root development upon phosphate deficiency and ammonium stress by modulating iron dynamics

**DOI:** 10.1101/2024.12.25.630297

**Authors:** Joaquín Clúa, Yves Poirier

## Abstract

Phosphorous and nitrogen are the main nutrients limiting plant growth in natural and agricultural systems. Plants typically uptake phosphorous in the form of inorganic phosphate, whereas the main nitrogen sources are nitrate and ammonium. Roots show a high degree of developmental plasticity in response to nutritional stresses, directly influencing plant fitness and stress resilience. In *Arabidopsis thaliana*, phosphate deficiency and high levels of ammonium in the media triggers a strong inhibition of the primary root growth. This developmental response is initiated by the accumulation of apoplastic iron in specific cell types, which triggers the inhibition of meristem maintenance and cell elongation. In this report, we show that DOMON domain-containing proteins are key molecular players controlling root development upon phosphate deficiency and ammonium toxicity via iron dynamics.

## Introduction

Phosphorous and nitrogen are the most important macronutrients for plant growth. Plants uptake phosphorous from the soil in the form of inorganic phosphate (Pi). Under natural and agricultural systems, Pi concentrations in the soil are typically low, limiting plant growth and crop production (Poirier *et al*., 2022). The main available forms of nitrogen for plants are nitrate (NO_3_^-^) and ammonium (NH_4_^+^). NH_4_^+^ is energetically more favorable since it is reduced and can be directly incorporated into amino acids and proteins. However, when NH_4_^+^ is present as the main source of nitrogen it triggers severe toxicity phenotypes (Li *et al*., 2014). Studies carried out in *Arabidopsis thaliana* have shown that Pi deficiency (-Pi) and NH_4_^+^ nutrition induce a strong inhibition of primary root growth (Britto & Kronzucker, 2002; Svistoonoff *et al*., 2007). Remarkably, these developmental responses are iron (Fe) dependent, as removal of Fe from the media or the addition of Fe chelators abolish root growth inhibition under -Pi and NH_4_^+^ nutrition (Svistoonoff *et al*., 2007; Liu *et al*., 2022b,a; Montpetit *et al*., 2022; Clúa *et al*., 2024). At the cellular level, it was recently revealed that root responses to both stresses are driven by tissue specific accumulation of apoplastic Fe (aFe), which in turn induces the arrest of meristematic activity and the inhibition of cell elongation (Müller *et al*., 2015; Balzergue *et al*., 2017; Gutiérrez-Alanís *et al*., 2017; Liu *et al*., 2022b; Naumann *et al*., 2022; Montpetit *et al*., 2022; Maniero *et al*., 2024).

At the molecular level, aFe dynamics depends on a tight regulation of Fe oxidation state (Liu *et al*., 2023; DeLoose *et al*., 2024). A high ratio of Fe_3_^+^/Fe_2_^+^ promotes aFe accumulation upon stress, which is dependent on the ferroxidase activity of LPR1 and LPR2. Loss-of-function mutations in LPR1 or LPR2 abolish both aFe accumulation and root growth inhibition under -Pi and NH_4_^+^ nutrition, respectively (Svistoonoff *et al*., 2007; Liu *et al*., 2022b; Naumann *et al*., 2022; Li *et al*., 2024). Recent studies showed that two members of the Cytochrome b561 and DOMON domain (CYBDOM) protein family, CRR and HYP1, are ferric reductases that play an antagonistic role to LPR1 under -Pi condition (Clúa *et al*., 2024; Maniero *et al*., 2024). Arabidopsis contains 10 CYBDOM proteins, which contain at least one N-terminal DOMON domain followed by a transmembrane cytochrome b561 domain. The CYBDOM proteins CRR and HYP1 are localized to the plasma membrane where they reduce aFe using cytosolic ascorbate as electron donor. Ectopic expression of CRR or HYP1 abolishes aFe accumulation due to a decrease in the Fe_3_^+^/Fe_2_^+^ ratio and abolishes the reduction of primary root growth under -Pi condition. Conversely, *crr* and *hyp1* mutants over-accumulate aFe and show a hypersensitive root phenotype under -Pi (Clúa *et al*., 2024; Maniero *et al*., 2024). These results suggest a molecular mechanism where the coordinated activity of LPRs and CYBDOMs proteins control root development under -Pi via aFe accumulation. However, whether CYBDOM proteins are also involved in root responses to NH_4_^+^ stress is unknown. Additionally, Arabidopsis genome contains a single gene, named AIR12, encoding a protein with a stand-alone DOMON domain containing a b-heme and capable of performing redox reactions with various substrates, such as superoxide anion radicals and monodehydroascorbate (Biniek *et al*., 2017). AIR12 is anchored to the plasma membrane via a GPI-tail and is facing the apoplast. Although AIR12 has been implicated in lateral root development, its role in the response of roots to -Pi and NH_4_^+^ stress as well as in aFe dynamics have not been investigated (Neuteboom *et al*., 1999; Preger *et al*., 2009; Costa *et al*., 2015; Gibson & Todd, 2015; Biniek *et al*., 2017; Wang *et al*., 2021).

## Results and discussion

To shed light on the function of *AIR12* under nutritional stresses and the potential redundancy with CYBDOM proteins, we analyzed primary root growth on +Pi and -Pi conditions of the *air12* single mutant relative to wild-type (Col-0), as well as to the *crr, hyp1*, and *crr hyp1* double mutant. Additionally, we generated a *crr hyp1 air12* triple mutant. Compared to Col-0, none of the mutants revealed defects in primary root growth under +Pi. As previously reported, *crr, hyp1*, and *crr hyp1* mutants were hypersensitive to -Pi, showing a short root phenotype (Clúa *et al*., 2024; Maniero *et al*., 2024). Interestingly, *air12* showed a significant reduction of about 18% in primary root elongation compared to Col-0, which is less severe than that of *crr* or *hyp1*. The introgression of *air12* in a *crr hyp1* mutant background did not have an impact on the double mutant phenotype (**Figure 1A, B**).

**Figure 1.**
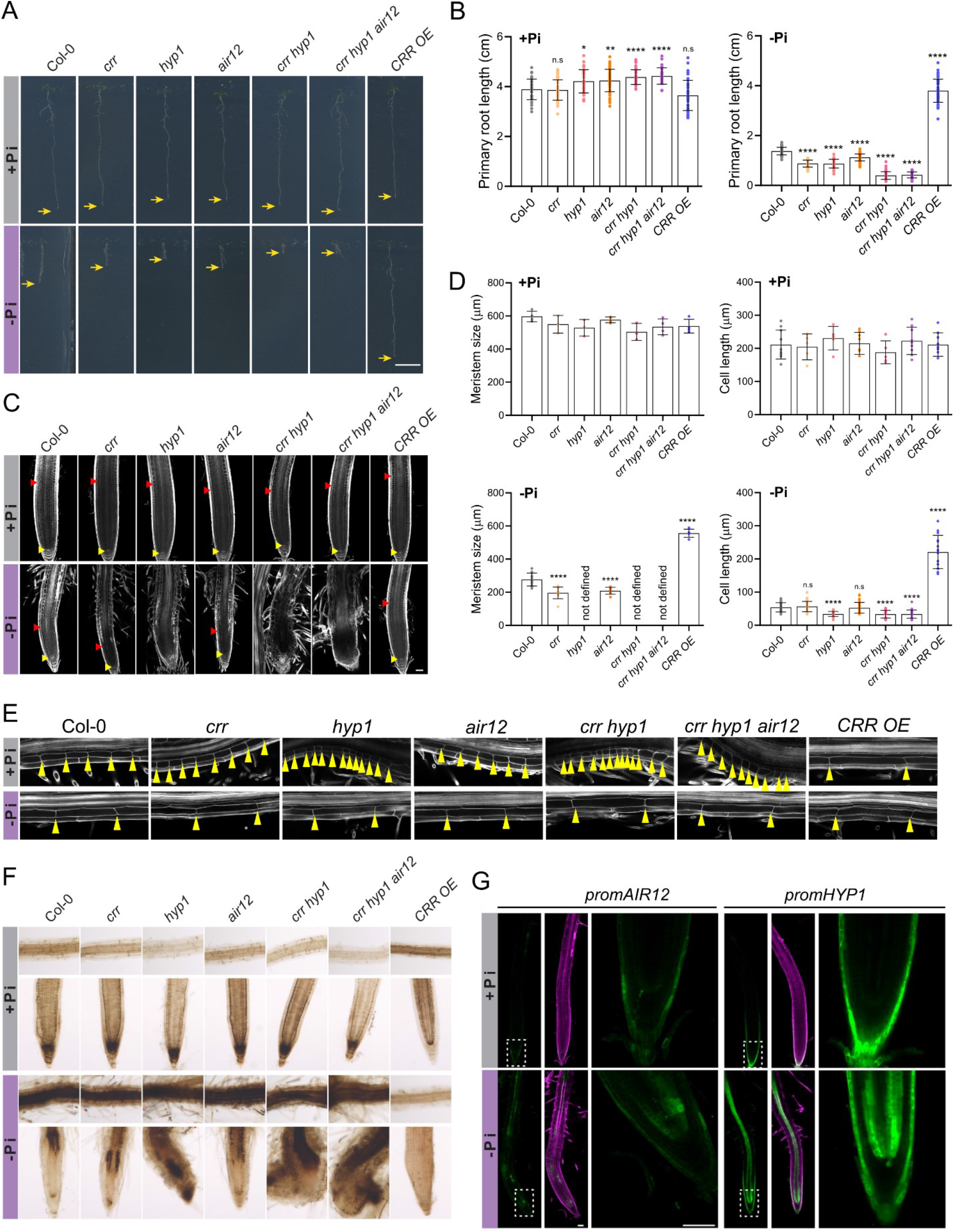
DOMON domain-containing proteins control root development under -Pi. **A**. Images showing root development under Pi sufficient (+Pi) or deficient (-Pi) conditions 7 days after germination. Yellow arrows indicate the root tip. Scale: 1 cm. **B**. Quantification of root length of the seedlings shown in A. **C**. Confocal images of root tips stained Calcofluor white. The yellow and red triangles indicate the quiescent center and the end of the meristematic zone, respectively. **D**. Quantification of meristem size and cell length in the differentiation zone of the root under +Pi and -Pi. In B and D bars show the mean and standard deviation. The statistical differences were analyzed using a one-way ANOVA followed by Dunnett’s multiple comparisons test. Significant differences to Col-0 are indicated (****P < 0.0001; ***P < 0.001; **P < 0.01; *P < 0.05). **E**. Confocal images showing the cell length in the differentiation zone. Yellow triangles indicate the boundaries between cells in the longitudinal axis. **F**. Visualization of the iron distribution using the Perls-DAB staining. The upper and lower panels show the root differentiation zone and root tip, respectively. G. Confocal microscopy images showing *HYP1* and *AIR12* expression pattern under +Pi and -Pi. The reporter lines were generated fusing HYP1 or AIR12 promoter to GFP (*promHYP1::GFP* and *promAIR12::GFP*). Cell walls were stained with Calcofluor white (magenta).

Analysis of root tips by confocal microscopy revealed that loss of *AIR12* further decreased meristem size under -Pi but did not exacerbate the inhibition of cell elongation showed by Col-0 (**Figure 1C-E**). Thus, *air12* mutant uncouples both growth processes similarly to *crr* (Clúa et al., 2024). In addition, both mutants show a reduction in the meristem size without altering its general structure or the shape and identity of its cells. This clearly contrasts with *hyp1, crr hyp1*, and *crr hyp1 air12* mutants, where the meristem lost its identity after 7 days post-germination (dpg) in -Pi (**Figure 1C**). Analysis of root tip morphology at different timepoints showed that these meristematic defects in the double and triple mutant are already evident at 4 dpg, showing aberrant cell proliferation (**Supplementary Figure 1**). Moreover, the microscopic analysis at different planes showed that from these undifferentiated mass of cells, new growth axes emerge (**Supplementary Videos 1 and 2**). Altogether, these data demonstrate that *AIR12* is required for meristem maintenance under –Pi condition and that the combined activity of *CRR* and *HYP1* is fundamental to coordinating cell divisions and cell shape upon -Pi.

To assess if the role of AIR12 in meristem maintenance under -Pi is also linked to aFe, we performed the Perls staining followed by DAB intensification to detect aFe in roots. Similarly to *crr, air12* mutant showed an increase of aFe accumulation under -Pi (**Figure 1F**). Thus, all the lines showed a positive correlation between aFe accumulation and the degree of root growth inhibition, further supporting the role of aFe in modulating the developmental response to -Pi. The addition of the Fe chelator ferrozine completely abolished the reduction of primary root growth observed in the *air12* mutant (**Supplementary Figure 2**).

We then analyzed the *AIR12* expression patterns and compared it to *HYP1* using transcriptional reporter lines (Costa *et al*., 2015; Maniero *et al*., 2024). In +Pi condition, *AIR12* is weakly expressed in the root cap, while under -Pi condition, the expression is further expended to the stem cell niche and vasculature (**Figure 1G**). This expression pattern of *AIR12* is similar to *HYP1* but weaker, and in both cases, it correlates well with the sites of aFe deposition. In conclusion, the data show that like CRR and HYP1, AIR12 contributes to primary root growth under -Pi condition and that its expression in the root meristem under -Pi is associated with a reduction of aFe in the meristem.

To investigate if DOMON domain-containing proteins play a role under NH_4_^+^ stress, we grew all mutant lines, as well as a line overexpressing CRR (CRR OE) (Clúa et al., 2024), in control conditions for 5 days followed by an additional 5 days on media containing either 5 mM of NO_3_^-^ or 5 mM NH_4_^+^ as the sole nitrogen source. Under NO_3_^-^ nutrition, none of the mutants or CRR OE showed strong differences in root elongation compared to Col-0 (**Figure 2A, B**). However, the quantification of root length under NH_4_^+^ nutrition reveals that *CRR* and *HYP1* are positive regulators of root growth under this condition. The *crr* mutant showed the strongest phenotype, with a reduction in root length of about 34% compared to Col-0, while *air12* did not show significant differences. The *crr hyp1* and *crr hyp1 air12* mutants showed a root phenotype similar to *crr*, suggesting that among the three DOMON domain-containing proteins, *CRR* plays the main role in NH_4_^+^ stress. Interestingly, the CRR OE line showed better root growth than Col-0, particularly under NH_4_^+^stress. This result indicates that manipulating the expression of CYBDOM proteins could be a valuable strategy to mitigate the root growth inhibitory effects of NH_4_^+^ (**Figure 2A, B**).

**Figure 2.**
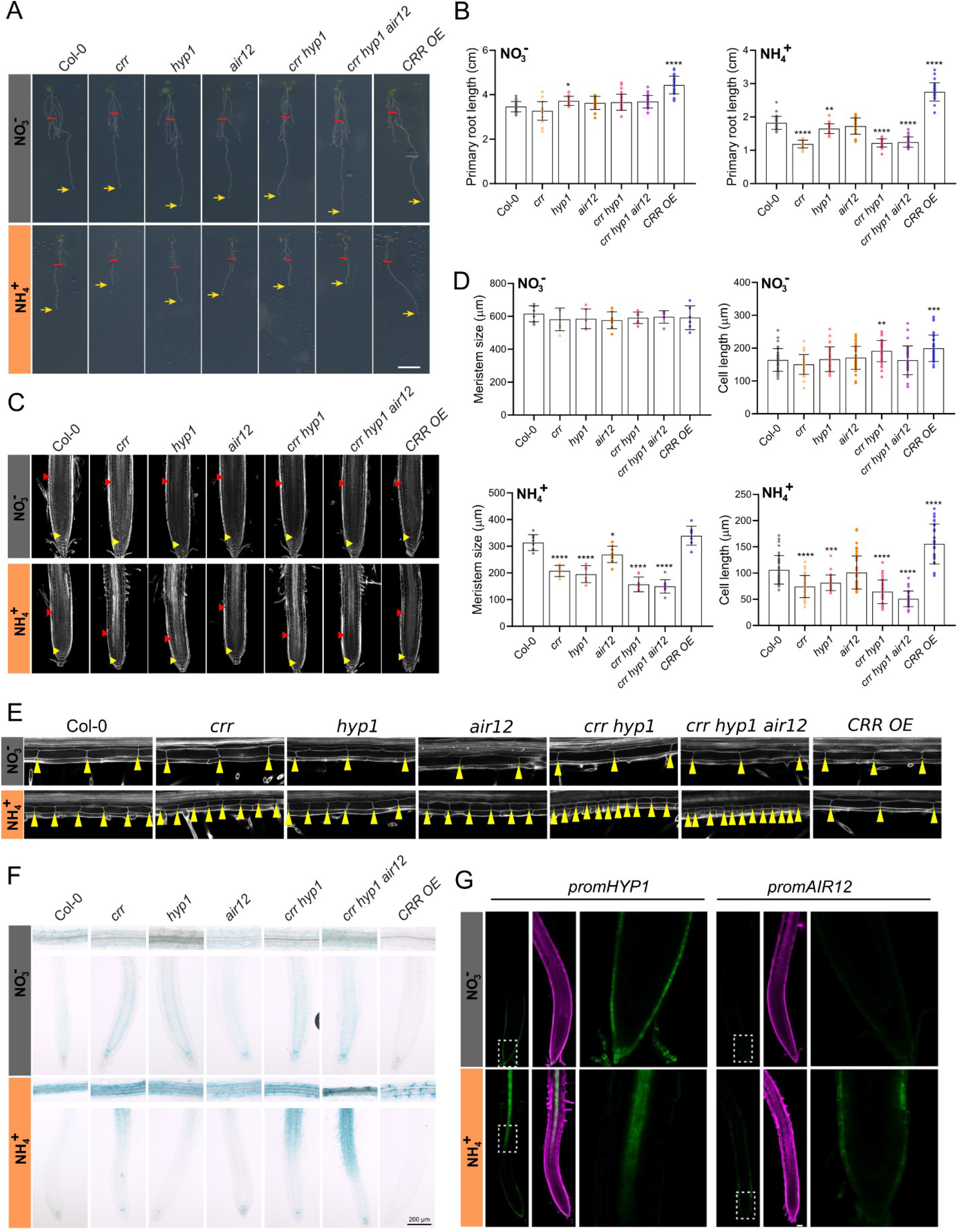
CRR and HYP1 are key determinants of ammonium toxicity resilience. **A**. Images showing root development under nitrate (NO_3_^-^) or ammonium (NH_4_^+^) nutrition. Seedlings were grown for 5 days in control conditions and then transferred to plates containing either NO_3_^-^ or NH_4_^+^ as the sole source of nitrogen for 5 additional days. Red lines and yellow arrows indicate the position of the root tip when the seedlings were transferred and at the end of the experiment, respectively. Scale: 1 cm. **B**. Quantification of root length of the seedlings shown in A. **C**. Confocal images of root tips stained Calcofluor white. The yellow and red triangles indicate the quiescent center and the end of the meristematic zone, respectively. **D**. Quantification of meristem size and cell length in the differentiation zone of the root under NO_3_^-^ or NH_4_^+^. In B and D bars show the mean and standard deviation. The statistical differences were analyzed using a one-way ANOVA followed by Dunnett’s multiple comparisons test. Significant differences to Col-0 are indicated (****P < 0.0001; ***P < 0.001; **P < 0.01; *P < 0.05). **E**. Confocal images showing the cell length in the differentiation zone. Yellow triangles indicate the boundaries between cells in the longitudinal axis. **F**. Visualization of the iron distribution using the Perls-DAB staining. G. Confocal microscopy images showing *HYP1* and *AIR12* expression pattern under NO_3_^-^ or NH_4_^+^. The reporter lines were generated fusing HYP1 or AIR12 promoter to GFP (*promHYP1::GFP* and *promAIR12::GFP*). Cell walls were stained with Calcofluor white (magenta).

Under NH_4_^+^ nutrition, Col-0 showed a reduction in both meristem size and cell elongation compared to the media containing NO_3_^-^ (**Figure 2C-E**). While the reduction in meristem size was further exacerbated in *crr, hyp1, crr hyp1* and *crr hyp1 air12*, CRR OE showed only marginal improvement in meristem maintenance compared to Col-0. In contrast to - Pi, the meristematic defects shown by *crr hyp1* and *crr hyp1 air12* mutants under NH_4_^+^ nutrition are relatively mild and have not produced aberrant cell divisions of altered cell shape. In terms of cell elongation, *CRR* and *HYP1* seem to play an additive role as the combined reduction in cell elongation of the single mutants is similar to the *crr hyp1* double mutant (40% reduction compared to Col-0, approximately). On the other hand, CRR OE dramatically reduced the inhibitory effect of NH_4_^+^ (**Figure 2D, E**). Interestingly, while CRR overexpression completely abolished the reduction of primary root growth observed under -Pi condition (**Figure 1B** and Clua et al. 2024), it did not completely abolish the reduction caused by NH_4_^+^ supply compared to the NO_3_^-^ control. This suggests that the modulation of the apoplastic Fe_3_^+^/Fe_2_^+^ ratio is not the only determinant of the root developmental response to NH_4_^+^. Further research will be needed to clarify which additional pathways contribute to root development under NH_4_^+^ nutrition.

Analysis of aFe distribution by Perls staining showed that Col-0 seedlings transferred to a media with NH_4_^+^ hyperaccumulate Fe in the differentiation zone of the root compared to NO_3_^-^ nutrition. This aFe accumulation pattern differs from that induced upon -Pi, which is characterized by strong aFe in the meristem. These distinct aFe signatures can explain the differences observed in the meristem structure of both stresses (**Figure 1F and 2F**). The *crr, crr hyp1* and *crr hyp1 air12* showed higher levels of aFe than Col-0 in the root elongation and differentiation zones. Additionally, in these mutants aFe accumulation was further extended to the meristematic zone, a region that was not affected in Col-0 upon NH_4_^+^ nutrition. In contrast, CRR OE showed an overall reduction in aFe independently of the nitrogen source (**Figure 2F**). The intensity of aFe deposition is thus positively correlated with root growth inhibition (**Figure 2A-B, F**). To further demonstrate the involvement of Fe dynamics in root responses to NH_4_^+^, we transferred Col-0, *crr hyp1, crr hyp1 air12*, and CRR OE lines to a media without Fe. Under these conditions, all the lines showed a similar root growth (**Supplementary Figure 3**), and the general toxic effects of NH_4_^+^ were greatly relieved (Liu *et al*., 2022b; Li *et al*., 2024).

Finally, we analyzed the expression pattern of *HYP1* and *AIR12*. Like in +Pi conditions, *HYP1* is mainly expressed in the root cap under NO_3_^-^ nutrition. However, upon NH_4_^+^ stress *HYP1* is additionally expressed in the root vasculature (**Figure 2G**). Interestingly, unlike under -Pi, *HYP1* is not detected in the stem cell niche upon NH_4_^+^ stress. This indicates that *HYP1* expression is tightly controlled in a tissue specific manner by factors which are stress-type dependent, and its expression is correlated with the aFe accumulation pattern (**Figure 1F, G and 2F, G**). On the other hand, we could only detect an induction of *AIR12* expression in the root cap under NH_4_^+^ nutrition. The fact that AIR12 expression is not detected in root zones where aFe normally accumulates upon NH_4_^+^ can explain the absence of phenotype of *air12* in this condition (**Figure 2A and 2G**). The endogenous expression pattern of *CRR* under NH_4_^+^ stress remains to be investigated as cloning of CRR with its promoter into T-DNA vectors fails to show adequate expression (Clúa *et al*., 2024). However, the analysis of publicly available single cell data shows that *CRR* is expressed in the meristem and in endodermal cells in control conditions (Clúa *et al*., 2024).

Altogether, our work shows that DOMON-containing proteins contribute to the adaptation of primary roots to -Pi and NH_4_^+^ stress. CRR, HYP1 and AIR12 differentially impacts growth and aFe accumulation under these distinct nutrient stresses, likely associated with their expression patterns in the root tip. We thus propose a model where DOMON domain-containing proteins, together with LPRs, constitutes an aFe-rheostat which adjust root growth upon both -Pi and NH_4_^+^ stress.

## Material and methods

### Plant material

All the mutants used in this work are in an *Arabidopsis thaliana* Col-0 background. The T-DNA insertion lines *crr-1* (Salk_202530) and *air12* (Salk_205490) were obtained from the European Arabidopsis Stock Center (Nottingham, UK). The *hyp1* (Salk_115548) T-DNA mutant was obtained from Ricardo Ghiel’s group (Maniero et al., 2024).

For the experiment related to Pi deficiency, the seeds were germinated on vertical square plates containing one-sixth-strength Murashige and Skoog (MS) salts without Pi (Duchefa Biochemie, Haarlem, the Netherlands), 1% (w/v) sucrose, 0.8% (w/v) agar (Plant Agar fromDuchefa Haarlem, the Netherlands), 0.5 gl^−1^ of 2-(N-morpholino)ethanesulfonic acid, and buffered to pH 5.8 with KOH. For the +Pi plates, the medium was supplemented with potassium phosphate buffer to obtain a final Pi concentration of 1mM. To simulate Fe deficiency in the ±Pi conditions a solution of ferrozine was added to the media to a final concentration of 100 µM.

For ammonium toxicity experiments, plants were grown on one-sixth-strength MS agar plates. After 5 days, seedlings were transferred to a media containing one-sixth-strength MS without nitrogen and supplemented with 5mM KNO_3_ or 2.5mM (NH_4_)_2_SO_4_ as the sole N source. In addition, plates were supplemented with Fe(III)NaEDTA to obtain the final Fe concentrations indicated. Otherwise stated, the final concentration of Fe used was 50 µM. Seedlings were photographed and analyzed 5 days after transfer.

### Generation of reporter lines

*AIR12* and *HYP1* promoter regions were selected based on previous reports (Costa et al., 2015; Maniero et al., 2024). PCR-generated promotor fragments were cloned into the pENTR2b (Thermo Fisher Scientific, Waltham, MA, USA) by Infusion, and recombined into the destination vector pFAST-G04 using Gateway technology (Shimada *et al*., 2010). Transgenic lines were generated via Agrobacterium tumefaciens-mediated transformations (Clough & Bent, 1998).

### Root length quantification

The plants were grown as described above and the plates were imaged with a flatbed scanner (Epson Perfection V700 photo; Epson, Suwa, Japan). The root length was measured using Fiji (http://fiji.sc/Fiji) and the plugin Simple neurite tracer.

### Determination of meristem size and cell length

The roots were treated with Clearsee and stained with Calcofluor White (Ursache *et al*., 2018), and observed using confocal microscopy. The average cell length of the cortical cells in the differentiation zone was calculated. The root apical meristem size was determined as the distance from the quiescent center to the first elongating cell.

### Histochemical detection of Fe in seedlings

Fe accumulation in seedlings was assayed using Perls staining as indicated (Meguro *et al*., 2003; Roschzttardtz *et al*., 2010). For Perls staining, the seedlings were incubated in 4ml of 2% (v/v) HCl and 2% (w/v) potassium ferrocyanide for 30min. The samples were washed with H_2_O and incubated for 45 min with 4ml of 10mM NaN_3_ and 0.3% H_2_O_2_ in methanol. After this step, the samples were rinsed with 100mM Na-phosphate buffer (pH 7.4) and were either analyzed or subjected to a DAB intensification step. For the latter, the seedlings were incubated for 30min in the same buffer containing 0.025% (w/v) DAB and 0.005% (v/v) H_2_O_2_ and washed two times with H_2_O. Finally, the samples were treated with chloral hydrate (1 gml^−1^; 15% glycerol) and analyzed with an optical microscope.

### Confocal microscopy

For confocal microscopy a Zeiss LSM 880 Airyscan (Carl Zeiss) microscope was utilized. Calcofluor White was excited at 405 nm and detected at 425–475 nm. GFP was excited with a 488 nm laser and the emitted light was collected at 493–538 nm. The analysis of confocal and bright field microscopy images was done with the ZEN 2 (blue edition) software from Zeiss and Fiji 2.9.0 (https://imagej.net/software/fiji/).

### Accession numbers

CRR, AT3G25290; AIR12, AT3G07390; HYP1, AT5G35735

## Acknowledgements

This work was supported by a grant from the Swiss National Science Foundation (31003A-182462) to Y.P.

