## Supplementary data for "DOMON domain-containing proteins control root development upon phosphate deficiency and ammonium stress by modulating iron dynamics"

**Supplementary Figure 1.** Kinetics of meristematic defects under -Pi

**Supplementary Figure 2.** Root responses to -Pi are iron dependent

**Supplementary Figure 3.** Root responses to $NH_4^+$ nutrition are iron dependent

**Supplementary Video 1.** Meristematic defects in *crr hyp1* double mutant

**Supplementary Video 2.** Meristematic defects in *crr hyp1 air12* triple mutant

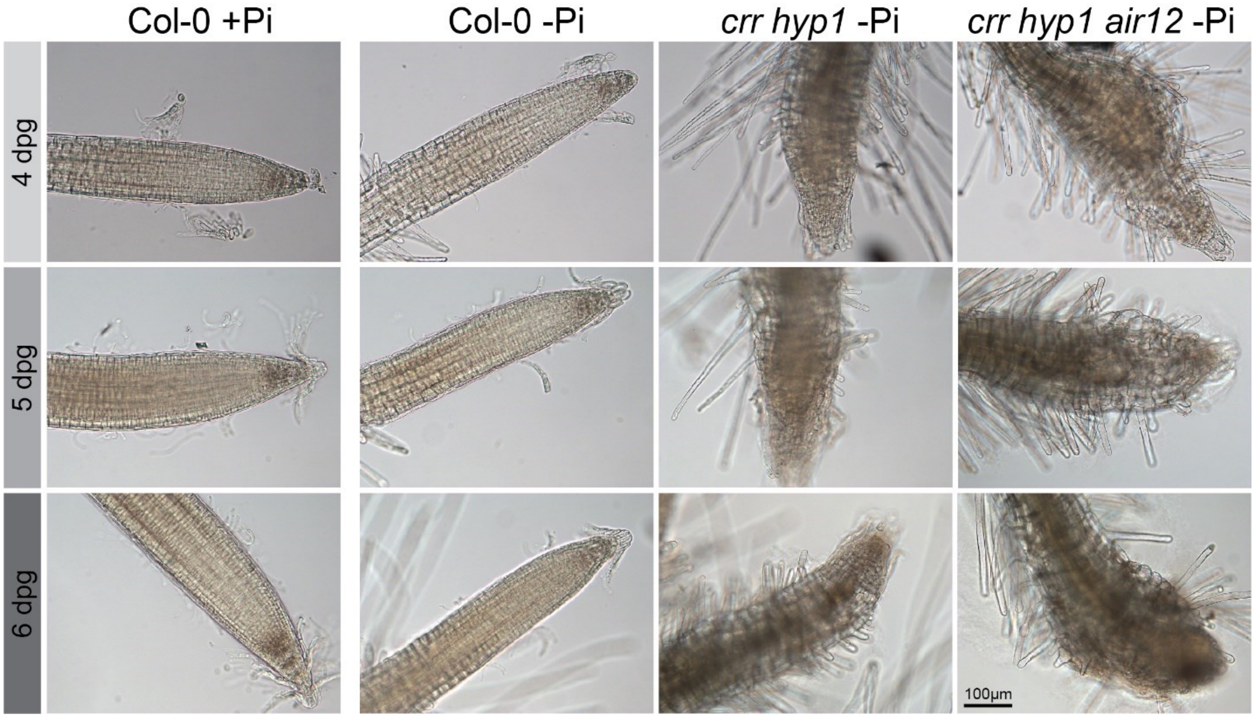


**Supplementary Figure 1. Kinetics of meristematic defects under -Pi**

Images showing the root apical meristem of plants grown under plus (+Pi) or minus (-Pi) phosphate conditions for the indicated amount of days after germination (dpg).

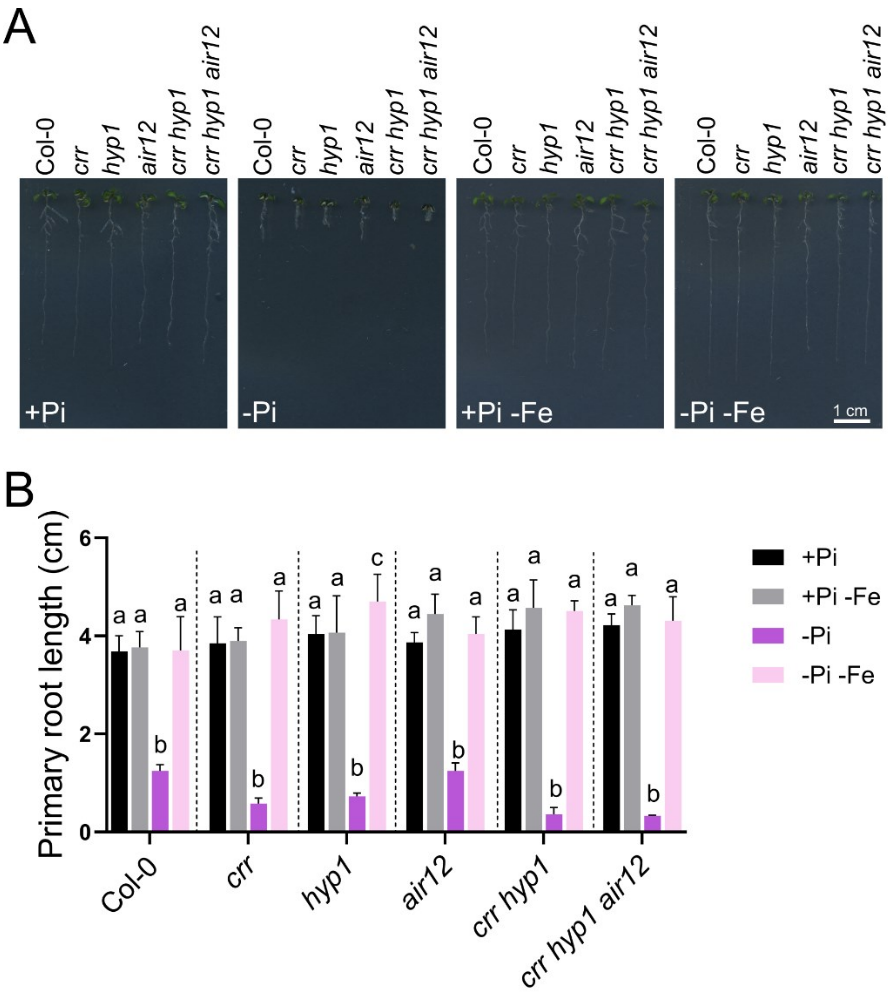


**Supplementary Figure 2. Root responses to -Pi are iron dependent**

**A**. Pictures showing the root phenotypes of plants grown for 7 days in media with or without phosphate (+Pi or -Pi, respectively). Ferrozine was added to the media to simulate iron deficiency (-Fe). **B**. Quantification of root length of seedlings grown in the indicated conditions. For each genotype, different letters indicate significant differences between growing conditions with a p-value< 0.05. Statistical differences were analyzed by a two-way ANOVA followed by a Tukey's multiple comparisons test.

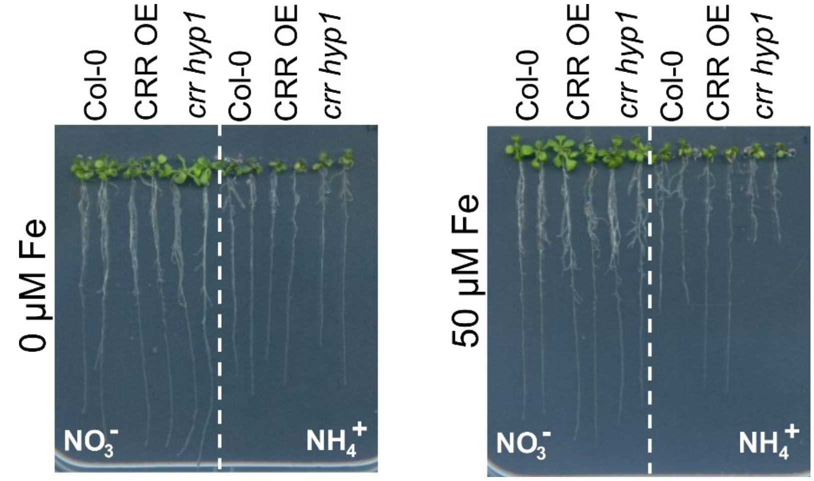


**Supplementary Figure 3. Root responses to $NH_4^+$ nutrition are iron dependent**

Root phenotypes of plants grown for 5 days in control conditions and then transferred to a media containing nitrate ($NO_3^-$) of ammonium ($NH_4^+$) as the sole source of nitrogen. The amount of additional iron added to the media is indicated.
